# Extremely thermophilic endospores germinate and metabolise organic carbon in sediments heated to above 80°C

**DOI:** 10.1101/2021.10.10.463673

**Authors:** Emma Bell, Jayne E. Rattray, Kathryn Sloan, Angela Sherry, Giovanni Pilloni, Casey R. J. Hubert

## Abstract

Endospores of thermophilic bacteria are widespread in cold seabed environments where they remain dormant during initial burial in accumulating sediments. The temperature increase during sedimentation can be simulated in experimental heating of sediments, resulting in the temperature-dependent activation of different endospore populations from the microbial seed bank. Here we investigated the response of endospore populations to heating at extreme high temperature (80– 99°C). Metabolites for germination and organic matter degradation (dipicolinic acid and organic acids) revealed both endospore germination and subsequent metabolism at ≥80°C. Endospore-forming *Firmicutes* with the genomic potential for organic carbon and nitrogen transformation were recovered by genome-resolved metagenomics. Genomes from *Symbiobacteriales, Thermosediminibacteriales, Moorellales* and *Calditerricolales* encode multiple mechanisms for high temperature degradation of sedimentary organic carbon and features of necromass that accumulate during sediment burial including saccharides, amino and nucleic acids. The results provide insight into the metabolism of novel carbon cycling microorganisms activated at high temperature, and suggest that extremely thermophilic *Firmicutes* dispersed in the ocean are poised to germinate in response to sediment heating during burial and transform a wide range of organic substrates.

## Introduction

Over one-third of marine sediments globally are estimated to be above 60°C and one-quarter above 80°C owing to subsurface thermal gradients (LaRowe et al., 2017). Microbial populations in these deeply buried sediments are therefore controlled by temperature and temperature-dependent physiological factors that the cells posess (Heuer et al., 2020). Sedimentation and burial of microorganisms deposited at the surface can take thousands of years (Kallmeyer et al., 2012). During burial cells remain viable (Orsi et al., 2013) and even metabolically active (Morono et al., 2020), but ongoing starvation in an environment gradually being depleted of available energy sources means that many microorganisms persist at very low energy yields or are dormant (Hoehler and Jørgensen, 2013; Jørgensen and Marshall, 2016). Persistence as endospores is one such form of dormancy that offers a survival strategy in energy-limited environments (Lomstein et al., 2012; Wörmer et al., 2019), potentially enabling populations adapted to warmer temperatures to persist during dispersal via burial through cooler, shallower intervals. Meanwhile, as the abundance and availability of organic carbon in sediments diminishes with depth owing to degradation by the activity of organotrophic communities nearer to the sediment surface, a decreasing fraction of organic matter remains in buried accumulated sediments (Middelburg, 2019). This leaves an altered and apparently diminished organic pool available for microbial populations in warmer deeper sediments.

Experiments with estuarine and deep subseafloor sediments designed to simulate sediment heating during burial show microbial acetate production, suggesting acetogenesis can provide fuel for microbial life in deeply buried anoxic sediments (Wellsbury et al., 1997, 2002; Parkes et al., 2007). Acetate is a key intermediate in the degradation of organic matter and can also be produced by CO_2_ reducing acetogens. Acetate provides both an electron donor for dissimilatory metabolism and a building block for biosynthesis, making it an important substrate in anoxic subsurface environments. Balanced microbial production and consumption of acetate in subsurface environments typically maintains acetate at a low concentration nearer to the surface (Wellsbury et al., 2002; Lever, 2012; Glombitza et al., 2019), whereas acetate accumulation has been reported in deeply buried anoxic sediments (Wellsbury et al., 1997; Egeberg and Barth, 1998; Heuer et al., 2020), mirroring sediment heating experiments that simulate burial (Wellsbury et al., 1997).

Sediment heating studies have demonstrated that endospores of thermophilic bacteria are widespread in cold surface environments where they cannot grow (Hubert et al., 2009; de Rezende et al., 2013; Müller et al., 2014; Volpi et al., 2017; Chakraborty et al., 2018; Hanson et al., 2019). Many of these studies have focused on sulfate-reducing bacteria, where acetate accumulation is expected from the incomplete oxidation of organic electron donors yielding acetate as a by-product (Widdel and Pfennig, 1977; Muyzer and Stams, 2008). Thermophilic endospores deposited in these cold seabed settings will remain dormant as they are buried by the accumulating sediments (de Rezende et al., 2013) and encounter heating during burial that could present an opportunity for temperature-dependent germination and growth of buried endospore populations (Hubert et al., 2010). In agreement with this, experimental heating of sediment applies a selective pressure that enriches distinct populations of endospore forming thermophiles at different temperatures (de Rezende et al., 2013; Bell et al., 2020). In those previous studies, endospore-forming sulfate reducers were actived upon heating to 50–70°C, whereas temperatures ≥80°C exceeded their thermal tolerance. This apparent cut-off has also been postulated for hydrocarbon biodegradation in sedimentary environments, as inferred from observations that oil biodegradation has not occurred in reservoir formations that have been buried to depths that are warmer than ∼80°C (Wilhelms et al., 2001; Head et al., 2003). However, it was recently demonstrated that the upper temperature limit for life in deeply buried marine sediments extends up to at least 120°C (Heuer et al., 2020). This means that populations in the hyperthermophilic temperature range are not necessarily restricted to mid-ocean ridge hydrothermal vents, where the highest known growth temperature have been recorded (Kashefi and Lovley, 2003; Takai et al., 2008). To explore the temperature-dependent germination and growth of endospore-forming bacteria in sediments at higher temperatures, we looked for metabolites of germinating endospores (dipicolinic acid) and organic matter degradation (organic acids) in incubations conducted at ≥80°C.

## Methods

### Sediment inoculum and medium preparation

Sediment collected from the Tyne estuary, United Kingdom (54°57’51’’N, 1°40’60’’W) was used as inoculum of thermophilic endospores in all sediment heating experiments. Anoxic sediment slurries were prepared by mixing sediment and anoxic seawater medium at a fixed ratio (1 g sediment to 2 mL medium) under a constant flow of N_2_ (Widdel and Bak, 1992; Isaksen et al., 1994).

### Temperature gradient incubations

Anoxic sediment slurries (10 mL) amended with lactate (final concentration 20 mM) were transferred to Balch tubes with a N_2_/CO_2_ headspace (1 bar pressure). Balch tubes were placed in a temperature gradient block at ∼5°C intervals from 80–99°C, with two replicates at each temperature. Heated anoxic sediment slurries were subsampled (2 mL) with an N_2_/CO_2_ flushed syringe after 24, 168 and 672 hours incubation. Subsamples were centrifuged (14,000 g, 5 min). The supernatant was used for measurement of free dipicolinic acid and organic acids. Subsamples for dipicolinic acid measurement were frozen -20°C then freeze-dried. Subsamples for organic acids were frozen and stored at -20°C.

### Dipicolinic acid (DPA) analysis

Endospores were quantified using the spore-specific biomarker 2,6-pyridine dicarboxylic acid (DPA). DPA dissolved in the medium i.e. ‘free DPA’ was used to quantify germinated endospores, as opposed to intact endospores where DPA remains in the spore core and requires release by hydrolysis (Rattray et al., 2021). Freeze-dried samples were re-constituted in 1 mL 1 M NaSO_4_, filtered and complexed with Tb^3+^. DPA was measured using HPLC fluorescence and DPA separation was performed using a Kinetex 2.6 μm EVO C18 100 Å LC column (150 × 4.5 mm, Phenomenex, USA) fitted with a guard column connected to a Thermo RS3000 pump. Gradient chromatography was used where solvent A was 1 M sodium acetate amended with 1 M acetic acid to pH 5.6 and solvent B was methanol/ultra-pure water (80:20). The sample injection volume was 50 μL and detection was perfomed using a Thermo FLD-3000RS fluorescence detector set at excitation wavelength 270 nm and emission 545 nm.

### Anoxic sediment enrichments

Anoxic sediment slurries (50 mL) were prepared in serum bottles with a N_2_ headspace. Two different carbon and energy sources were tested: (1) sediment-derived organic matter already present with no extra substrate addition; (2) amendment with a complex substrate mixture of tryptic soy broth (3 g/L), glucose (3 mM) and the carboxylic acids acetate, propionate, butyrate and lactate (3 mM each). Heated anoxic sediment slurries were subsampled (1.5 mL) with an N_2_ flushed syringe after 113 and 279 hours incubation. Subsamples were centrifuged (13,000 g, 5 min) with the resulting supernatant used to measure organic acids and the sediment pellet used for DNA extraction.

### Organic acid measurements

Organic acid measurements from temperature gradient incubations were performed at the University of Calgary, Canada, where formate, acetate, propionate, lactate, butyrate and succinate were measured using UV (210 nM) on an HPLC RSLC Ultimate 3000 with an Aminex HPX-87H, 7.8 × 300 mm analytical column. The HPLC flow rate 0.6 mL/min and the eluent was 5 mM H_2_SO_4_. Organic acid measurements from sediment enrichments were performed at Newcastle University, UK, where acetate, propionate and butyrate were measured by ion (exclusion) chromatography (IC) using a Dionex ICS-1000 with an AS40 auto-sampler equipped with an IonPac ICE-AS1, 4 × 250 mm analytical column. The IC flow rate was 0.16 mL/min, the eluent was 1.0 mM heptafluorobutyric acid and the cation regenerant solution used for the AMMS-ICE II Supressor was 5 mM tetrabutylammonium hydroxide.

### DNA extraction and 16S rRNA gene amplicon sequencing

DNA was extracted from sediment pellets using the PowerSoil DNA isolation Kit (MoBio Laboratories) following the manufacturer’s protocol with minor amendments (Bell et al., 2018). Extracted DNA was used as a template for PCR amplification using Golay barcoded fusion primers targeting the V4-V5 region of the 16S rRNA gene (Caporaso et al., 2012). PCR reaction components and cycling conditions are described in the study by Bell *et al*., (2018). PCR products derived from a common sub-sampling time from triplicate sediment slurries were in most instances pooled prior to clean-up using Agencourt Ampure XP paramagnetic beads resulting in a single pooled amplicon library for a given experimental time point. 16S rRNA gene amplicons were sequenced on an Ion Torrent Personal Genome Machine (School of Natural and Environmental Sciences, Newcastle University, UK) in accordance with the manufacturer’s instructions (Life Technologies). Sequencing data were processed by the Torrent Suite Software V4.0. Raw sequence reads were demultiplexed and quality filtered in QIIME version 1.9.1 (Caporaso et al., 2010). All subsequent sequence analysis was performed with USEARCH v11 (Edgar, 2010). Sequences were truncated to 350 bp (*fastx_truncate*) and clustered into operational taxonomic units (OTUs) sharing 97% sequence identity with UPARSE (*cluster_otus*) (Edgar, 2013). Taxonomy was predicted with SINTAX (Edgar, 2016) with a USEARCH compatible (Lee, 2020) Silva 138 database (Quast et al., 2013). Amplicon data were visualised with the R package Ampvis2 (Andersen et al., 2018). Normalised OTU counts were used to calculate the correlations between each OTU and the concentration of organic acids at the corresponding sampling time points using the Python library Pandas (The Pandas Development Team, 2021).

### Metagenomic sequencing, assembly, binning and analyses

Metagenomic sequencing was performed on an Illumina NovaSeq 6000 with a S4 300 cycle flow cell. Libraries were prepared by shearing to an insert size of ∼200 bp using a Covaris instrument, followed by library construction with the NEB Ultra II DNA library prep kit. Reads were preprocessed with BBDuk (Bushnell et al., 2017) and assembled separately with two assemblers (1) metaSPAdes (Nurk et al., 2017) and (2) MEGAHIT (Li et al., 2015) using the *meta-sensitive* option. Raw reads were mapped to each of the assemblies with BBMap (Bushnell et al., 2017). Each of the assemblies were binned with both MetaBAT2 (Kang et al., 2019) and CONCOCT (Alneberg et al., 2014). Bins from the same assembler were refined using DAS Tool (Sieber et al., 2018). The best bins from each of the approaches were selected with dRep (Olm et al., 2017) using the parameters; completeness 75%, contamination 5%, primary cluster average nucleotide identity (ANI) 90%, secondary cluster ANI 99%. This resulted in a total of 13 representative metagenome assembled genomes (MAGs).

Protein-coding genes were predicted with prodigal (Hyatt et al., 2010). Amino acid sequences used to predict the optimum growth temperature of microorganisms using Tome (Li et al., 2019). MAGs from microorganisms predicted to be thermophiles were annotated with METABOLIC v4.0 (Zhou et al., 2019), which integrates annotation of protein-coding genes with KOfam (Aramaki et al., 2020), TIGRfam (Selengut et al., 2007), Pfam (Finn et al., 2014), dbCAN2 (Zhang et al., 2018) and MEROPS (Rawlings et al., 2016). 16S rRNA gene sequences in the metagenomic dataset were identified with METAXA2 (Bengtsson-Palme et al., 2015) and phyloFlash (Gruber-Vodicka et al., 2020). 16S rRNA sequences from the metagenomic dataset were aligned to 16S rRNA gene amplicon sequences with BLAST (Altschul et al., 1990).

MAGs were taxonomically classified with GTDB-Tk v1.5.0 with reference data for GTDB R06-RS202 (Chaumeil et al., 2019). A phylogenomic tree of thermophile MAGs was constructed with GToTree v1.5.39 using the *Firmicutes* specific single-copy gene HMM profile (119 genes) (Lee, 2019). To create the tree, genomes from the four families (*ZC4RG38, Calditerricolaceae, Thermosediminibacteraceae* and *Moorellaceae*) were downloaded from GTDB release 95 using *gtt-get-accessions-from-GTDB* in GToTree.

## Results

### Metabolites show endospore germination and activity at high temperature

Endospores contain ∼25% of a 1:1 chelate of DPA with divalent cations, predominantly Ca^2+^ (Anil et al., 2008). DPA gets rapidly released when spores initiate germination (Setlow, 2006), resulting in an extracellular pool of ‘free DPA’ (Rattray et al., 2021) that was measured in temperature gradient incubations of estuarine sediment. Free DPA accumulated after 24 h incubation at 80– 99°C, indicating that endospores present in the sediment germinated upon heating resulting in DPA release into the culture medium. Using a spore-specific DPA conversion factor that accounts for the higher DPA content of thermophilic spores (Rattray et al., 2021) it was estimated that between 1.3–2.1 × 10^4^ endospores mL^-1^ germinated in the slurries within 24 h incubation at 80–99°C (Fig. 1A). A small concentration of formate (<0.5 mM) was detected at all temperatures after 24 h (Dataset S1) with additional accumulation of formate, acetate and propionate after 168 h (Fig. 1B) indicating further hyperthermophilic metabolism by the germinated bacteria at these high temperatures. Lactate was the only organic carbon source added to the temperature gradient incubations, and was not consumed at any temperature (Dataset S1) suggesting that accumulation of these metabolites is due to degradation of sedimentary organic matter (total organic carbon 6.4 ± 0.1 %; (Bell et al., 2018)). At the highest temperature (99°C) fewer spores germinated after 24 hours (Fig 1A), presumably owing to temperature limitations of corresponding vegetative cells. Accordingly, organic acid production at 99°C was comparably lower and exhibited greater variability between replicates (Fig. 1B). The accumulation of free DPA in the sediment after 168 and 672 h suggests continued germination.

**Figure 1:**
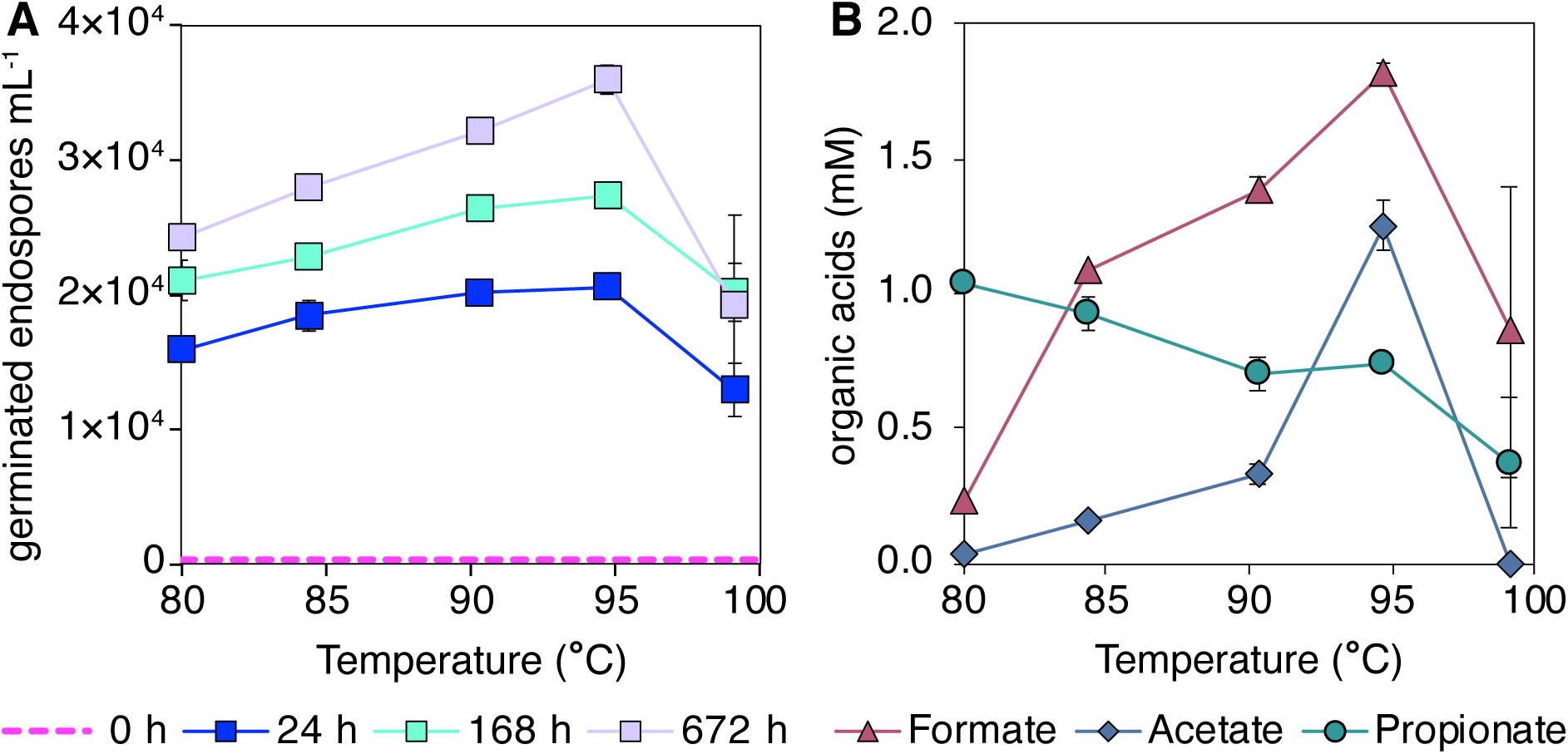
Metabolites associated with endospore germination (free DPA) and organic matter degradation (organic acids) in heated sediment. (A) Germinated endospore counts determined from the free DPA concentration in estuarine sediment incubated at 80–99°C in a temperature gradient block for 24, 168 and 672 hours. The baseline (0 h) is the average of two replicate DPA measurements from unincubated sediment. (B) Formate, acetate and propionate production in estuarine sediment incubated at 80–99°C in a temperature gradient block for 168 hours. Error bars show standard deviation of two replicate incubations. Raw data is provided in Dataset S1.

### High read abundance of *Firmicutes* in thermophilic sediment enrichments

To assess endospore forming bacteria in the hyperthermophilic temperature range in greater detail, larger sediment slurries were heated to 80 and 90°C (Fig. 2A) and DNA was extracted after 113 and 279 h incubation. Acetate and propionate production were observed (Fig. 2A), similar to the temperature gradient incubations with the same sediment (Fig. 1B) (formate was not quantified for the larger sediment slurries). Addition of organic carbon resulted in elevated concentrations of organic acids compared to the parallel unamended sediment incubations, with acetate and propionate reaching 8 and 2 mM, respectively (Fig. 2B). Tracking 16S rRNA gene amplicon read abundance of endospore-forming *Firmicutes* showed a marked increase following sediment heating (Fig. 2C), consistent with endospores in the sediment germinating upon incubation at high temperature. A small number of OTUs accounted for the majority of 16S rRNA gene amplicon reads, with *Symbiobacterales* and *Thermosediminibacter* increasing to the greatest relative abundance in 80 and 90°C incubations (Fig. 2C). *Thermanaeromonas* and *Caldicoprobacter* OTUs were also detected, but only at 80°C (Fig. 2C), suggesting 90°C exceeds the growth limit in some cases. Pearson correlation of each OTU with organic acid concentration showed high correlation (>0.6) for *Thermosediminibacter* and *Thermanaeromonas* OTUs (Fig. S1).

**Figure 2:**
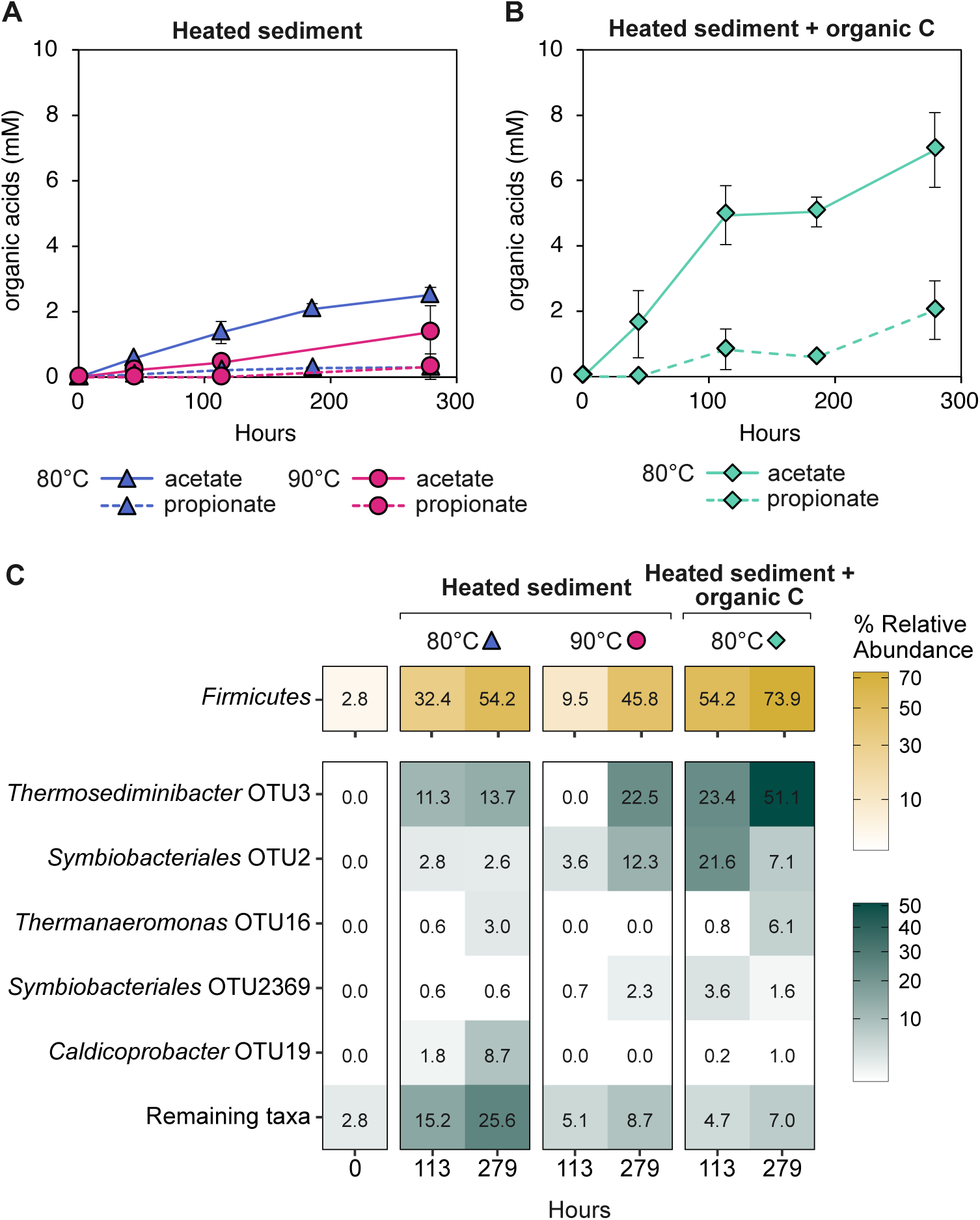
Organic acid production and enrichment of *Firmicutes* in sediment heated to 80 and 90°C. (A) Acetate (solid line) and propionate (dashed line) production in sediment heated to 80°C (blue triangles) and 90°C (pink circles). (B) Acetate (solid line) and propionate (dashed line) production in sediment heated to 80°C and supplemented with organic carbon (green diamonds). Error bars show standard deviation of three replicate incubations. (C) Corresponding 16S rRNA gene amplicon relative abundances from heated anoxic sediments enrichments after 113 or 279 hours of heating. The top bar (yellow shading) shows the read abundance assigned to the endospore-forming phylum *Firmicutes*. The bottom panel (green shading) shows *Firmicutes* OTUs with the greatest read abundance in heated sediment incubations. Read abundances for all OTUs are provided in Supplementary Dataset S3.

### Genome-resolved metagenomics applied to heated sediments

To explore the metabolic potential of endospore populations enriched at high temperature two sediment slurry samples incubated at 80°C were selected for metagenomic sequencing. These samples were selected to capture populations revealed by amplicon sequencing to be present at both 80 and 90°C (Fig. 2C). Assembled metagenomic contigs were binned and dereplicated resulting in 13 MAGs from four phyla (*Firmicutes, Proteobacteria, Actinobacteriota* and *Campylobacterota*; genome completeness is provided in Dataset S4). Optimum growth temperatures for each of the thirteen MAGs was predicted based on inferred amino acid composition (Li et al., 2019) and used to predict which organisms could be metabolically active at high temperature (Dataset S4). Four MAGs affiliated with the phylum *Firmicutes* were predicted to be thermophiles with growth temperature optima from 69–75°C (Fig. 3), consistent with the activity observed in the high temperature sediment incubations. The other nine MAGs were predicted to be mesophiles with temperature optima ranging from 24–29°C, and were likely binned from relic DNA (Lennon et al., 2018) arising from organisms that were abundant *in situ* and died at elevated temperature.

**Figure 3:**
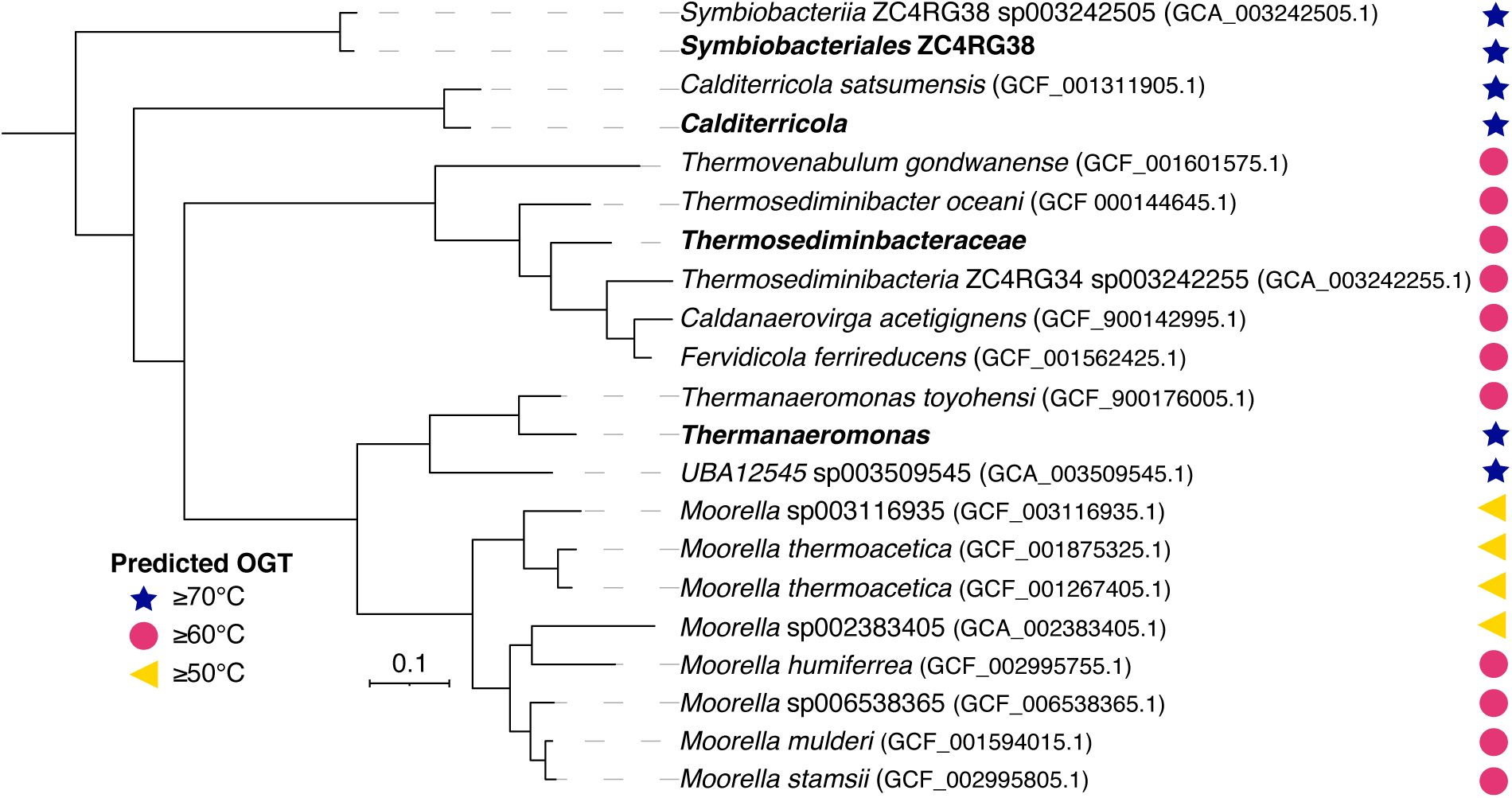
Phylogenomic tree with predicted optimum growth temperature (OGT) of thermophilic endospore-forming *Firmicutes*. Four high quality MAGs from this study are shown in bold. Representative genomes from the same families (*ZC4RG38, Calditerricolaceae, Thermosediminibacteraceae* and *Moorellaceae*) were downloaded from GTDB and included in the tree. The tree is based on 119 concatenated single copy genes. The scale bar corresponds to per cent average amino acid substitution over the alignment and supports that all four genomes represent novel lineages within their respective phylogenetic groups. Amino acid based predictions of OGT were made using Tome (Li et al., 2019).

Phylogenomic analysis of the *Firmicutes* MAGs revealed taxonomic affiliations with: (1) the family ZC4RG38 (class *Symbiobacteriales*); (2) the family *Thermosediminibacteraceae*; (3) the genus *Thermanaeromonas*; and (4) the genus *Calditerricola* (Fig. 3). *Symbiobacteriales, Thermosediminibacteraceae* and *Thermanaeromonas* MAGs all contained 16S rRNA gene sequences that were ≥99% identical to the amplicon library OTUs with high read abundances (OTUs 2, 3 and 16; Fig. 2C). An unbinned *Calditerricola* 16S rRNA gene sequence was recovered from the metagenome that was 99.7% identical to a low read abundance *Calditerricola* amplicon (0.02%; OTU 783) and is presumed to belong to the same population, since no other *Calditerricola* 16S rRNA gene sequences were recovered from the amplicon or metagenomic datasets. Comparision of these four extreme thermophile MAGs to genomes in the Genome Taxonomy Database (GTDB R06-RS202) did not uncover any close relatives (>95% AAI). The amino acid identity (AAI) to the closest relatives ranged between 79.9–93.3% (Fig. 3) indicating that these MAGs represent new genera and species within their respective phylogenetic groups (Jain et al., 2018). Temperature optima predictions for genomes from organisms in the same family, showed that these related genomes also belong to thermophiles (shown in the phylogenomic tree in Fig. 3). Accordingly, these related organisms were discovered in geothermal subsurface aquifers (Mori et al., 2002; Ogg and Patel, 2009) and high temperature compost (Moriya et al., 2011; Martins et al., 2013).

### Genomic evidence for sporulation and dormancy in extreme thermophiles

Thermophilic *Firmicutes* enriched from estuarine sediments are predicted to be spore-formers, based on DPA release in temperature gradient incubations of the same sediment (Fig. 1A), and their viable persistence in an environment much below their temperature requirement for growth. Consistent with this, all extreme thermophile MAGs contained core sporulation genes that are conserved in well-known spore-forming *Bacilli* and *Clostridia* (Galperin et al., 2012). These include genes required for pre-septation (Stage 0), post-septation (Stage II), post-engulfment (Stages III-VI), spore coat assembly and germination (Dataset S5). Notably, the four MAGs also include genes for DPA synthesis (*dapG, asd, dapA* and *dpaAB*) and SpoVA proteins that create a channel for DPA movement across the spore inner membrane during sporulation and that facilitate its release as free DPA during germination (Christie and Setlow, 2020).

### Metabolic potential for organic carbon and nitrogen degradation

The build up of organic acids in unamended sediment heating experiments indicates that extreme thermophiles degrade sedimentary organic matter as well as glucose and tryptic soy compounds provided in the organic carbon amendment. Multiple pathways for acetate production from organic compounds were present in *Symbiobacteriales* and *Thermosediminibacteraceae* genomes (Fig. 4). Both possess the phosphoenolpyruvate (PEP):carbohydrate phosphotransferase system (PTS) for uptake and concomitant phosphorylation of carbohydrates (Deutscher et al., 2006). This includes enzyme I (encoded by the *ptsI* gene) and HPr (encoded by the *ptsH* gene) as well as a sugar-specific enzyme II (EII) permease. Multiple EII permeases and ABC transporters were found (Fig. 4) including those for the uptake of plant and algal derived saccharides (e.g., cellobiose and mannibiose). Both genomes contained multiple glycoside hydrolases (Fig. 4) for the breakdown the glycosidic bonds in carbohydrates producing free fermentable glucose. *Symbiobacteriales* can subsequently produce acetate via ADP-forming acetyl-CoA synthetase (*acdAB*) while *Thermosediminibacteraceae* can produce acetate in a two-step conversion via phosphate acetyltransferase (*pta*) and acetate kinase (*ackA*). Both organisms encode enzymes that couple formate oxidation to hydrogen evolution, *Symbiobacteriales* via a membrane bound [NiFe] Group 4a-hydrogenase, and *Thermosediminibacteraceae* via a cytosolic [FeFe] Group A4-hydrogenase (Søndergaard et al., 2016). *Thermanaeromonas* and *Calditerricola* do not harbour genes for acetate production and are therefore unlikely contributors to the acetate production observed in heated sediment incubations. Instead, the presence of AMP-forming acetyl-CoA synthetase (*acs*; 6.2.1.1) in both *Calditerricola* and *Thermanaeromonas* genomes suggest these organisms are acetate consumers.

**Figure 4:**
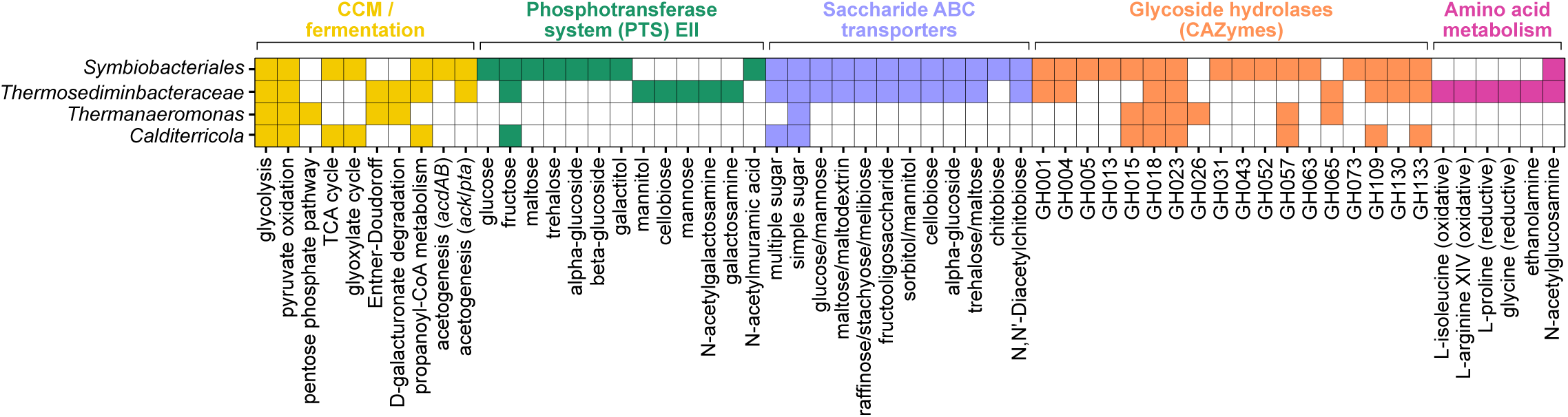
Metabolic potential of extremely thermophilic *Firmicutes*. Presence of select carbon and nitrogen metabolising pathways in four high-quality MAGs of thermophilic endospore-forming *Firmicutes*. Details of KEGG modules, CAZymes and MetaCyc pathways used to create the figure are provided in Dataset S6.

Necromass from cells that died at elevated temperature offer another source energy in these sediment heating experiments. Cellular necromass contains mostly proteins and amino acids as well as DNA, RNA and membrane sugars (Orsi et al., 2020). *Thermosediminibacteraceae* has the genomic potential to co-ferment amino acids via Stickland reactions, using isoleucine and/or arginine as electron donor and proline and/or glycine as electron acceptor (Fig. 4). This MAG can also degrade ethanolamine, a compound readily available from the degradation of cell membranes. Both *Thermosediminibacterace* and *Symbiobacteriales* have EII permeases for uptake of amino sugars (e.g., *N*-acetyl glucosamine and glucosamine). Amino sugars phosphorylated by the PTS system enter glycolysis following the removal of acetyl and amino groups by the enzymes *N*-acetylgluscosamine-6-phosphate deacetylase (*nagA*) and glucosamine-6-phosphate deaminase (*nagB*). Purines from dead cells offer another source of organic carbon and nitrogen as *Thermanaeromonas* had the potential to degrade xanthine, a nitrogen-rich organic compound from nucleic acids that is widespread in aquatic environments (Cunliffe, 2016). *Thermosediminibacteraceae, Symbiobacteriales* and *Thermoanaeromonas* genomes all contained an extracellular nuclease for degradation of DNA polymers and all MAGs contained nucleoside and nucleobase transporters that may be used for salvage or as a carbon and energy source (Pérez Castro et al., 2021).

### Metabolism of sulfur compounds

Sulfate reduction does not occur in these sediments when incubated at ≥80°C (Bell et al., 2020). *Thermanaeromonas* has genes for dissimilatory sulfite reductase (*dsrAB*), the sulfur relay protein *dsrC* and the electron transport complex *dsrMKJOP*. The absence of sulfate adenylyltransferase (*sat*) and adenylylsulfate reductase (*aprAB*) in this high quality MAG suggests that this organism may instead utilise sulfite as an electron acceptor. The closely related isolate *T. toyohensis* is also unable to respire sulfate (Mori et al., 2002). *Thermanaeromonas* could potentially yield sulfite from thiosulfate (*phsA*) or from the desulfonation of sulfolactate by sulfolactate sulfo-lyase (*syuAB*), that cleaves *R*-sulfolactate into pyruvate and sulfite (Denger and Cook, 2010). Sulfolactate is a widespread natural product in plants, algae and prokaryotes, and is also a component of bacterial endospores that gets released upon germination (Bonsen et al., 1969; Rein et al., 2005). Sulfolactate produced by germinating endospores could therefore provide a source of organosulfate in heated sediments. Sulfolactate and free DPA release during spore germination could be providing additional nourishment and nutrients to thermophiles following their activation at high temperature.

## Discussion

High temperature incubation of surface sediment resulted in the germination and activity of extremely thermophilic endospore populations that did not reduce sulfate (Dataset S1). In these sediments, the activity of sulfate reducing bacteria is limited to <80°C (Bell et al., 2020), in line with the maximum reported growth temperature among endospore forming sulfate reducers being 85°C (Nazina et al., 1988). In sediments heated to these temperatures, extremely thermophilic organotrophs germinate and metabolise different pools of organic carbon including sedimentary organic matter, resulting in the production of organic acids and the accumulation of acetate. Sedimentary organic carbon typically contains 10–20% carbohydrates, 10% nitrogenous compounds and 5–15% lipids with the remaining fractions consisting of unidentified organic compounds that have been labelled recalcitrant (Arndt et al., 2013). *Symbiobacteriales* and *Thermosediminibacteraceae* both encode multiple pathways for degradation of plant and algal derived saccharides, as well as amino sugars from chitin, fungal cell walls and bacterial outer membranes. These organisms accounted for the greatest proportion of reads in both unamended and organic carbon amended incubations (Fig. 2), and appear to be responsible for the accumulation of organic acids in the heated sediments.

Endospore formation and widespread dispersal of microorganisms in aquatic environments (Müller et al., 2014; Mestre and Höfer, 2021) means extremely thermophilic *Firmicutes* have the potential to be transported long distances and dispersed to seafloor sediments worldwide. *Thermosediminibacter oceani*, the closest relative to the *Thermosediminibacteraceae* detected in this study (Fig. 3), was isolated from sediments 1–2 metres below seafloor (mbsf) (Lee et al., 2005) where the *in situ* temperature only 12°C, whereas *T. oceani* grows optimally at 68°C. Although spore formation was not reported for this strain, this highlights the ability of closely related thermophiles in deeply buried sediments to become active in response to heating. Upon germination, this organism can metabolise amino acids which are the most abundant fermentable organic matter in deep subseafloor sediments (Lomstein et al., 2012; Orsi et al., 2020).

Earlier studies that report acetate production in heated estuarine and deep subseafloor sediments (≤90°C) attribute this observation to the temperature activation of organic carbon increasing its bioavailibilty, thus providing a continuous supply of energy for bacteria and archaea during burial in the deep biosphere (Wellsbury et al., 1997; Parkes et al., 2007). Recent perspectives on organic matter degradation in the marine biosphere present a more complex understanding, suggesting that organic matter reactivity is ecosystem dependent (LaRowe et al., 2020; Dittmar et al., 2021). This implies microbial, geochemical and environmental factors affect the potential for metabolism such that recalcitrance is not necessarily an intrinsic property of a given compound (Middelburg, 2018). This more nuanced perspective is consistent with a role for temperature activation of dormant populations of thermophilic endospores capable of organic matter degradation, and supports earlier suggestions that changes in the reactivity of organic carbon with temperature is due to changes in the prevailing metaproteome, i.e., by the activation of dormant thermophiles and expression of their enzymes (Hubert et al., 2010).

Sediments in this study were heated to ≥80°C for the duration of the incubation period. These temperatures are commonly used for pasteurisation, where it is expected that vegetative cells will be killed after exposure for up to 1 hour. Dead cells offer an abundant source of proteins, nucleic acids, polysaccharides and lipids. Compounds released by germinating endospores could similarly provide sources of carbon and energy in these heated sediments. Sulfolactate (5% dry weight) and DPA (5–15% dry weight) are biodegradable compounds released during germination (Bonsen et al., 1969; Banerji and Regmi, 1998; Setlow, 2006; McClintock et al., 2018). *Thermanaeromonas* was the only spore-former detected in this study with the genomic potential to metabolise sulfolactate, whereas all MAGs associated with endospore-forming bacteria contained genes for DPA uptake. The anaerobic transformation of DPA to acetate, propionate, ammonia and CO_2_ has been reported for a coculture of two marine microorganisms (Seyfried and Schink, 1990), but exact enzymatic pathways required for DPA fermentation have not been elucidated (Gupta et al., 2019). Nevertheless, the presence of DPA in the sediment after 672 h incubation does not suggest the microbial degradation of DPA by these extreme thermophiles.

Surface sediments used in these and other heating experiments contain more organic carbon than would be encountered in deeply buried marine sediments. Nevertheless they offer insight into the metabolic potential of extremely thermophilic organotrophs that can potentially survive as spores during burial and germinate upon heating in a deeper hotter environment where necromass and metabolites released during germination could support their organotrophic metabolism as vegetative cells. This approach remains a useful model for predicting microbial ecology and organic geochemistry in the deep subsurface, where biomass levels are too low to facilitate direct metagenomic studies (Dombrowski et al., 2018; Heuer et al., 2020). Shallow sediments seed deeper sediments (Inagaki et al., 2015), where community assembly is driven by selection mechanisms that filter out populations with unfavourable traits (Petro et al., 2017). The resilience of endospores makes them suited to not being filtered out, possibly explaining estimates that endospores account for a significant proportion of microbial biomass in deeper sediments where they have been proposed to outnumber vegetative cells (Lomstein et al., 2012; Wörmer et al., 2019). Deep sediments thus harbour a seed bank of endospores with genomic and functional diversity that can remain viable for long periods of time, enabling their selection when environmental circumstances are appropriate. Following a long journey of dispersal, any of the thermophilic endospore populations detected and described here could germinate to become active members of the hot deep biosphere by metabolising deeply buried organic matter.

## Data availability

Data analysed in this study are provided in the supplementary information. Sequence data have been deposited in GenBank under the BioProject PRJNA371432.

## Acknowledgments

This work was supported by a UK Natural Environment Research Council award to CRJH (NE/J024325/1) and EB (NE/K501025/1). Research grants to CRJH from the UK Engineering and Physical Sciences Research Council (EP/J002259/1) and ExxonMobil Research and Engineering (New Jersey) through the Knowledge Build program, and by a Campus Alberta Innovates Program (CAIP) chair awarded to CRJH.

---

**Figure S1:**
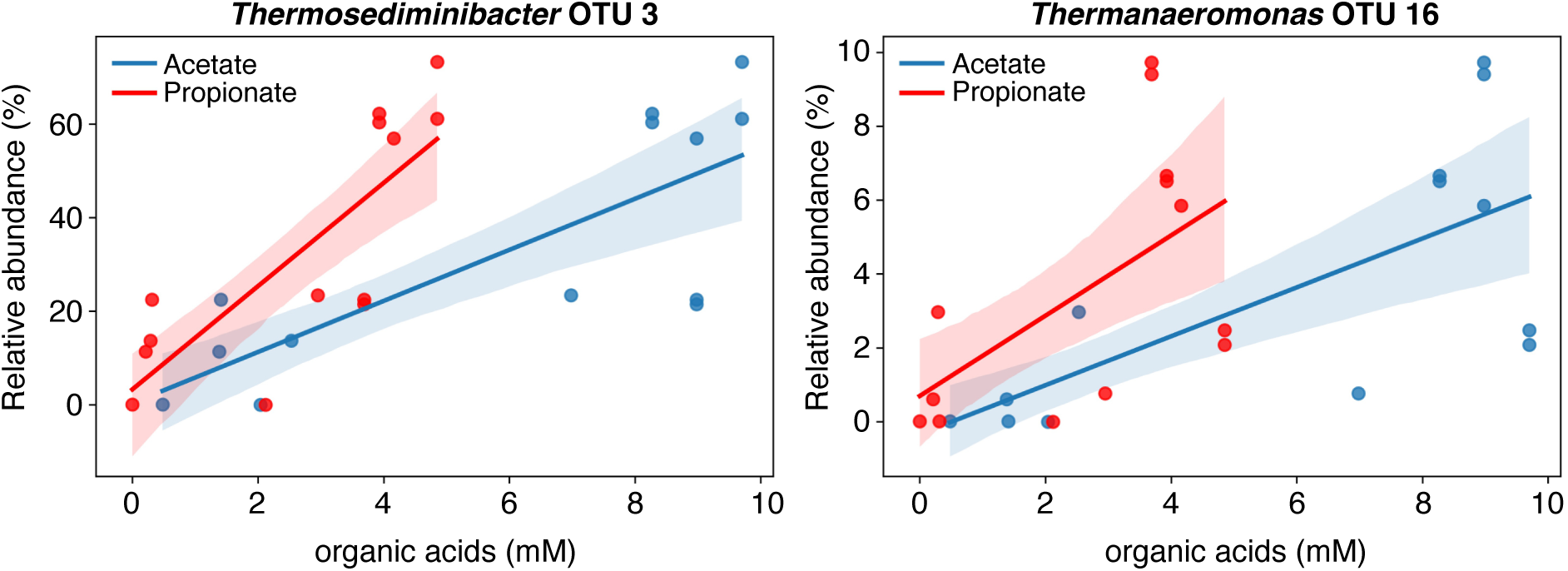
Relative abundance select OTUs plotted against the organic acid concentration at the correspondent sampling time points. The two most abundant OTUs with high correlation to acetate (Pearson correlation >0.6) are shown.

